# Standardised TruAI automated quantification of intracellular neuromelanin granules in human brain tissue sections

**DOI:** 10.1101/2025.01.20.633975

**Authors:** Anastasia Filimontseva, Thais Cuadros, Zac Chatterton, Amy Burke, Anahid Ansari Mahabadian, Joan Compte, YuHong Fu, Miquel Vila, Glenda M Halliday

## Abstract

**Aims:** To standardise and automate the quantitation of human-unique neuromelanin granules in catecholamine neurons in post-mortem tissue sections from healthy individuals at different ages to understand any changes in these granules with age.

**Methods:** 5-6 µm-thick fixed and paraffin-embedded transverse midbrain tissue sections were supplied from 47 cases from three brain banks following ethical approvals. Sections were prepared and automated digital images acquired. Standardisation and automation of the quantification of neuromelanin granules was performed using the TruAI feature of the Olympus VS200 desktop platform. Comparisons between stained and unstained sections as well as correlations with age were performed.

**Results:** The automated platform reliably identified both stained and unstained intracellular neuromelanin granules and extracellular pigments, showing high reproducibility in measurements across laboratories using different tissue processing methods. Extraneuronal pigments were significantly smaller than intracellular neuromelanin granules. Sections processed for haematoxylin and eosin staining impacted the size and colour of both neuromelanin and the neurons containing neuromelanin. Haematoxylin made neuromelanin bluer, and the increased tissue processing made the intracellular area occupied by neuromelanin smaller in younger people. There was an increase in neuromelanin optical density and colour change (browner) with age.

**Conclusions:** The TruAI automated platform reliably quantifies individual neuromelanin granules in catecholamine neurons. Extraneuronal pigments are considerably smaller in size than intracellular neuromelanin, and intracellular neuromelanin changes its properties with age. The darkening and colour change of intracellular neuromelanin suggests an increase in eumelanin over time in healthy individuals. These changes can be reliably identified using the automated platform.

**Key points:**

- Standardised, reliable TruAI automated quantitation of intracellular neuromelanin granules in human brain sections gives novel insights into their structure and function
- Extraneuronal pigments, including neuromelanin granules released from dying pigmented neurons, are significantly smaller than intracellular neuromelanin granules
- Intracellular neuromelanin granules change their properties with healthy ageing becoming darker, consistent with more antioxidant eumelanin

## Introduction

Artificial intelligence (AI) is now frequently used to objectively analyse diverse images and has transformed many medical analyses by providing diagnostic and pathological classifications [1-3]. Such AI platforms provide more efficient and objective outcomes to the traditional manual scoring or counting methods previously used, saving time and improving accuracy in classifications and other analyses. Many public bioimaging repositories now have open-source semi-automated AI quantification platforms and workflows readily available [4-6]. Despite these advances, there are limitations in these quantification workflows for particular types of analyses. For example, some automated analyses are only applicable to small regions of interest (ROIs) [1]. This is inconvenient because whole-slide/image analyses are necessary for understanding pathological disease mechanisms, especially in neurodegenerative research. Additionally, publicly shared deep learning algorithms are streamlined for overall general use rather than assessment of specialised cellular organelles. This maximises reproducibility across diverse types of studies, but user-friendly protocols for the analysis of these more specific cellular features also need development.

Human catecholaminergic neurons contain a unique intracellular pigment – neuromelanin granules – which begin to accumulate at around 3-5 years of age and are fully developed between 20 and 50 years of age [7]. These neurons are selectively vulnerable to Parkinson’s disease (PD), the second most common neurodegenerative disorder that manifests in motor and non-motor symptoms usually in late adulthood [8]. These symptoms arise due to gradient depigmentation and loss of catecholaminergic neurons in the substantia nigra pars compacta (SNpc), locus coeruleus and ventral tegmental area [8, 9]. Cell vulnerability is associated with neuromelanin accumulation, but the exact contribution of this macromolecule in PD remains unclear due to the time-consuming quantitation required across several anatomical regions in large-sized human post-mortem sections, and the lack of endogenous neuromelanin in catecholaminergic neurons in rodent or primate models. Despite this, the role of neuromelanin in cell pathogenesis has been investigated at the regional level by quantifying the number of pigmented cells in catecholaminergic regions [8, 10] and more recently by measuring the optical density of intracellular neuromelanin granules (iNM) [11-14]. However, due to different histological preparations of sections and the use of different optical density measurements, iNM analyses have not been standardised, and a high-capability analysis platform is needed for more robust, reproducible and efficient analyses.

We present an automated image acquisition and quantification protocol for iNM analyses in human post-mortem sections. The platform can be used on digital images of unstained or haematoxylin and eosin (H&E) stained tissue by applying the AI-trained automated quantification of neuromelanin using the Olympus VS200 desktop technology (EVIDENT Technology GmbH, ver. 4.1.1 build 29408). The platform allows the quantification of the size, optical density and intracellular area occupied by iNM in these histological preparations, and also the quantification of previously unexplored red, green and blue colour features of iNM. This is the first platform for the automated standardized analysis of iNM in human brain sections and it has revealed surprising significant differences in the size of neuromelanin granules as well as identified potential histological processing effects that impact on age associations.

## Materials and Methods

The platform and workflow are presented in Figure 1. Briefly, tissue is processed and formalin-fixed paraffin-embedded sections are prepared for H&E staining or are kept unstained. For automated digital image acquisition, we used the SLIDEVIEW VS200 Slide Scanner (Olympus Corporation, Tokyo, Japan). To automate the quantification of iNM, we used the TruAI™ feature of the Olympus VS200 desktop technology (EVIDENT Technology GmbH, ver. 4.1.1 build 29408, https://evidentscientific.com/en/software).

**Figure 1.**
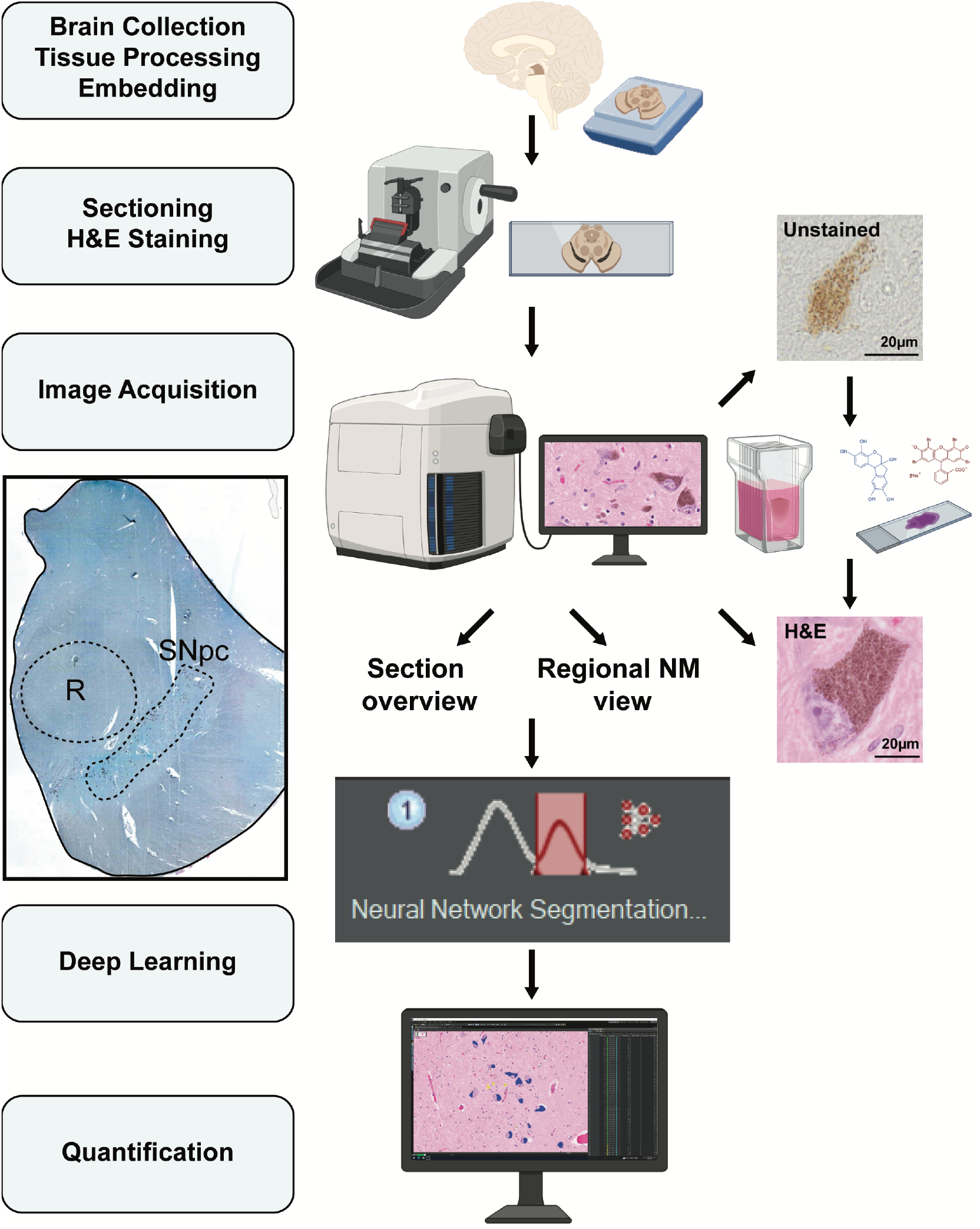
Platform and workflow for neuromelanin granule (NM) automatic quantification. Brain samples are collected and undergo standard processing procedures. After sectioning, images are acquired by scanning the slides. Anatomical delineations of neuromelanin-containing regions (such as the substantia nigra pars compacta, SNpc) and nearby landmarks (such as red nucleus, R) are identified. Lastly, standardized features of intracellular neuromelanin are determined by applying the AI trained automated quantification protocol.

### Case selection and tissue processing

Five to six µm transverse brainstem sections through the midbrain at the level of the red nucleus were provided by three ethically approved brain banks (see ethics approvals) from 21 healthy control cases aged from 21 to 100 years old (used for healthy age related studies) and 26 additional cases overlapping in age but with neurological disease (used for maximizing and validating the cutoff for extracellular neuromelanin from dying cells). Case information and section use are given in the table in the supporting information. Sections were deparaffinized by incubation at 60°C for 30 min to 1 hr.

To assess unstained neuromelanin granules, sections were submerged in HistoChoice Clearing Agent (Sigma-Aldrich, H2779) for 2 × 7 min, followed by rehydration in decreasing ethanol concentrations (100% ethanol for 2 × 3 min, 95% ethanol for 3 min, 70% ethanol for 3 min) and distilled H2O for 3 min. Slides were submerged in phosphate-buffered saline and coverslipped using a Fluorescence Mounting Medium (DAKO, S302380-2) for imaging.

To assess H&E stained iNM granules, sections were submerged in xylene for 3 × 3 min and rehydrated in decreasing ethanol concentrations (100% ethanol for 2 × 5 min, 95% ethanol for 2 × 5 min, 70% ethanol for 2 × 5 min) and double distilled H2O for 5 min. Sections were stained with Harris’ Haematoxylin solution for 2 min and rinsed for 5 min under running tap water, then 0.1% Eosin for 2 min, washed by dipping in water, and differentiated in 70% ethanol for 3 min. Sections were dehydrated in increasing ethanol concentrations (96% ethanol for 3 min, 100% ethanol for 2 × 5 min) xylene for 2 × 5 min. Sections were mounted in DPX organic mounting media.

To assess cresyl violet-stained iNM granules, sections were submerged in xylene for 3 × 3 min and rehydrated in decreasing ethanol concentrations (100% ethanol for 2 × 5 min, 95% ethanol for 2 × 5 min, 70% ethanol for 2 × 5 min) and double-distilled H2O for 5 min. Sections were stained with 0.5% cresyl violet for 2 min, washed by dipping in water, and dehydrated in 100% ethanol for 2 × 3 min then xylene for 2 × 3 min. Sections were mounted in DPX organic mounting media.

### Image acquisition

Digital images using the SLIDEVIEW VS200 automated slide scanner (Olympus Corporation, Tokyo, Japan) with pre-set focusing and exposure parameters. Brightfield images were captured using a 20x X Line™ extended apochromat objective (UPLXAPO20X), which has an extended range of chromatic aberration compensation from 400 nm to 1000 nm. A focus map was generated for each sample using the semi-automatic autofocus function to optimise scan speed. Images were captured as a single image z-stack or an extended focal imaging scan, which merges multiple z-planes into a single sharp image (see detailed scanning protocol in the Supplementary materials).

### TruAI software training

The software was trained to identify unstained and H&E-stained neuromelanin automatically. For both section types, background regions and NM-positive areas were manually annotated using the brush tool across 5–10 scanned sections (see Figure 2). Consistent training labels were applied across images to ensure standardisation (see Figure 2), and these labelled images were exported as inputs for model training (see detailed scanning protocol in the Supplementary materials).

**Figure 2.**
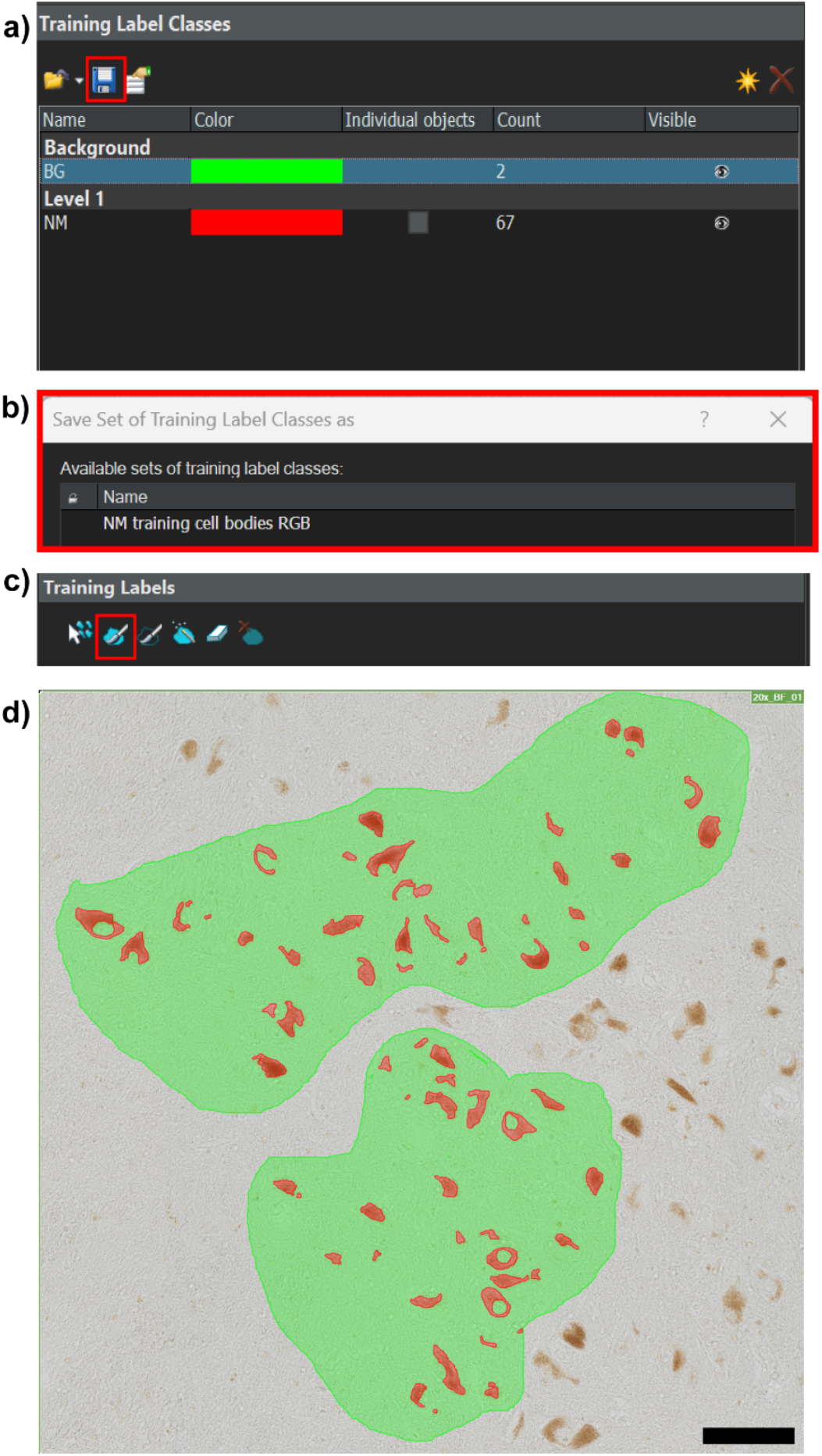
Training labels for neuromelanin (NM) quantification. a) Creation of a new training label class for NM granules. b) This training label must be saved and applied to sections. c) NM granules need to be accurately traced with the brush tool. d) Annotations need to be applied to 5-10 sections. Scale bar equals 100 μm.

### TruAI deep learning

Once at least five sections are annotated, the deep learning training can begin. Training label images were loaded into the program and used to generate a neural network model. The required accuracy for neural training is a minimum of 0.85 similarity. For the first attempt, ‘no limit’ can be selected under training duration to allow for any modifications during the neural training. To increase the accuracy of neural training, it is recommended that more sections be annotated. Once the training reached the desired threshold, the checkpoint with the highest similarity score was selected and saved. This trained model was then used for all future NM quantification (see detailed scanning protocol in the Supplementary materials).

### Automated TruAI quantification

The trained neural network algorithm was applied to all scanned images for automated quantification of neuromelanin. ROIs were manually drawn on both unstained and H&E-stained images using the “Count and Measure” tool. The trained model was then applied to these ROIs to detect NM (see Figure 3). Individual NM objects were highlighted by the software, and the results appeared in the “Count and Measure Results” window. Relevant parameters such as area and mean RGB intensity were selected for analysis of iNM (see Figure 3), with optional size thresholding applied prior to export (see detailed scanning protocol in the Supplementary materials).

**Figure 3.**
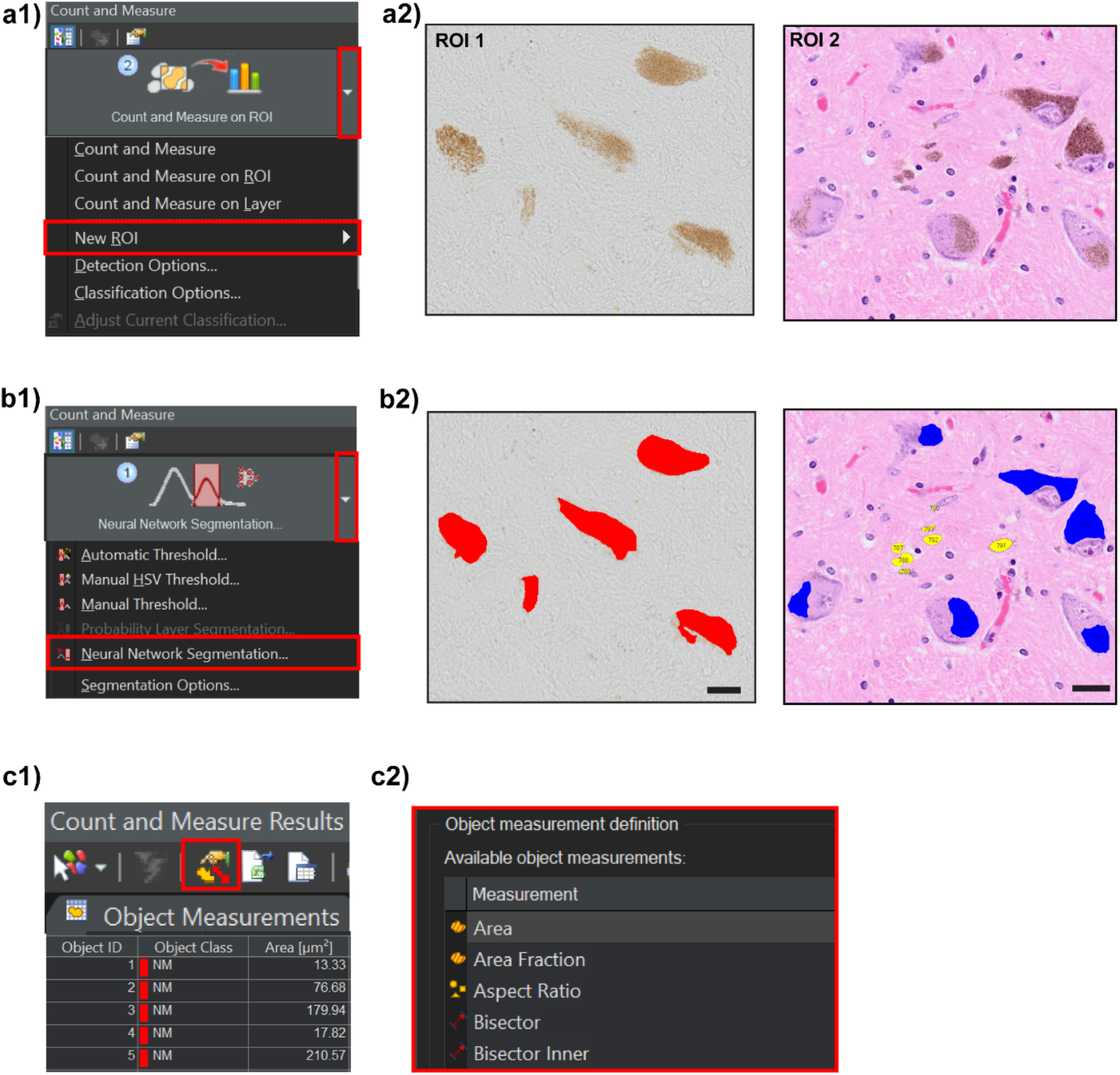
Automated TruAI quantification of neuromelanin. a1, a2) In selected sections, a new ROI needs to be drawn. b1, b2) The automated quantification of neuromelanin will run on the drawn ROI using the ‘Neural Network Segmentation’ option. Scale bars equal 20μm. c1, c2) A variety of features can be explored after completing neural network segmentation.

### Statistical analysis

Statistical analyses were performed using R, IBM SPSS (SPSS Inc., Chicago, IL, USA, version 26) and GraphPad Prism (GraphPad Software Inc., California, USA, version 10.0.3) statistical analysis software. Statistical significance was set at *P* < 0.05. For neuromelanin granule size thresholding, K-means clustering (k=2) was performed to determine size thresholds for intracellular versus extraneuronal pigments. Scripts used for the analysis are available from https://github.com/zchatt/Neuromelanin. For intracellular neuromelanin, Mann-Whitney tests were performed to determine differences between stained and unstained sections in the area occupied by neuromelanin, the estimated grey optical density, and the colour intensity, with linear regression used to assess age related changes.

## Results

### The size of intracellular neuromelanin granules versus extraneuronal pigments

Analysis of the size of iNM and extraneuronal pigments revealed that extraneuronal pigments were smaller than iNM granules. TruAI automated analysis was performed on unstained neuromelanin granules and their size examined in different substantia nigra pars compacta dopamine neuron subclusters, including: the ventral tier (SNV), lateral part (SNL) and dorsal tier (SND). The distribution of neuromelanin granule size (log10) in SNV, SNL and SND revealed two distinct modes (Figure 4a). K-means clustering was performed to identify iNM and extraneuronal pigment size thresholds based on these two modes. The threshold for iNM granules was above 78.3µm^2^, and the extraneuronal pigment cutoff was below 58.5µm^2^ (Figure 4b). Scripts for this analysis are available from https://github.com/zchatt/Neuromelanin/tree/1.0.

**Figure 4.**
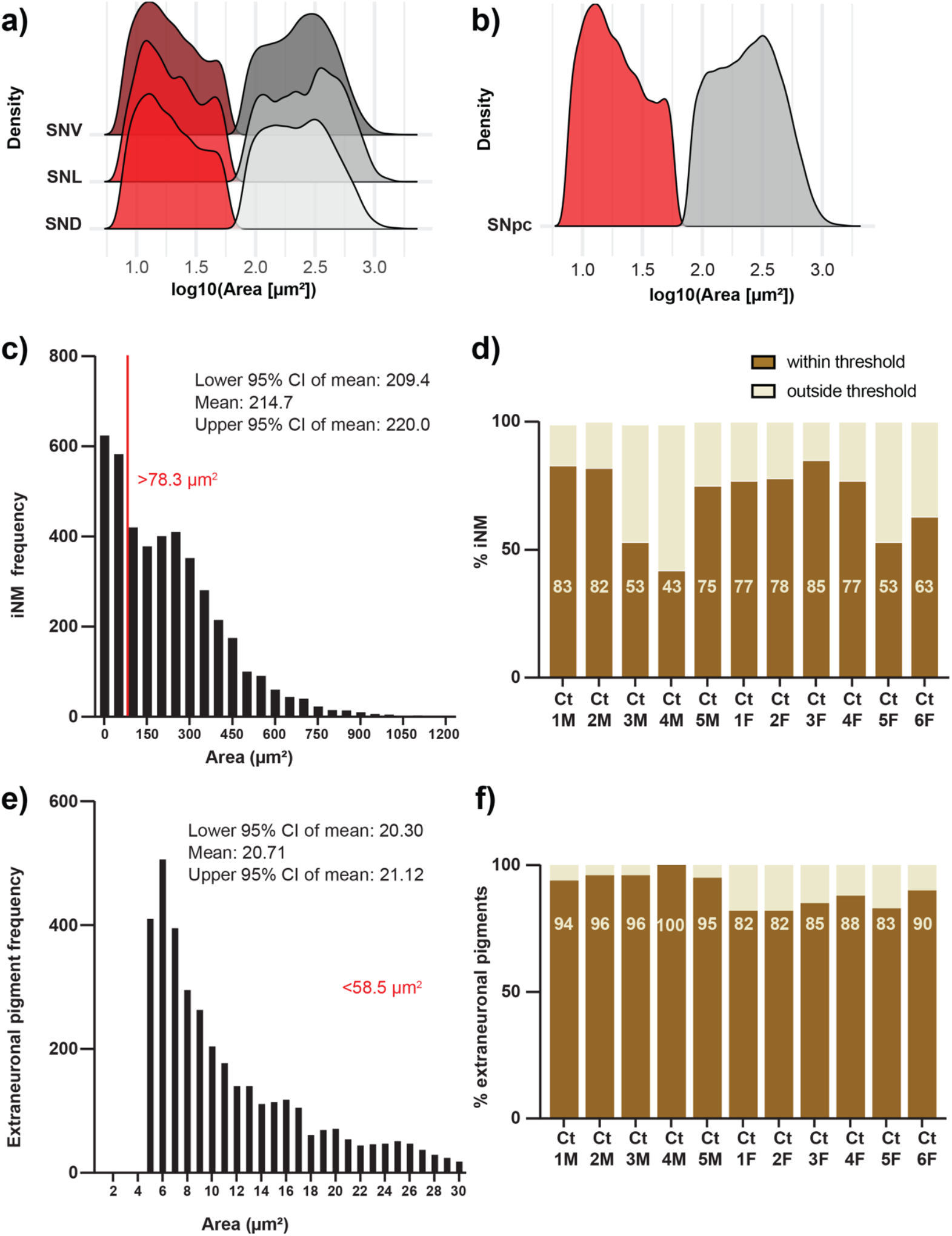
Neuromelanin size thresholding. a) Distribution of neuromelanin size (log10) measured in unstained sections of the substantia nigra pars compacta ventral tier (SNV), lateral part (SNL), and dorsal tier (SND). K-means clustering was performed in each subregion independently. This defined extraneuronal pigments as <58.5µm^2^ (red histogram) and iNM as >78.3µm^2^ (grey histogram). b) Combined neuromelanin granule size distribution across the substantia nigra pars compacta (SNpc). c) iNM size threshold applied to identified H&E stained iNM granules. The distribution of iNM size with respect to the defined threshold is shown. d) Proportion of identified H&E stained iNM granules included within the defined threshold. e) Extraneuronal pigment area threshold applied to identified H&E stained extraneuronal pigments. f) Proportion of identified H&E stained extraneuronal pigments included within the threshold.

This classification model was validated using H&E-stained sections. By using the iNM granule size threshold, we observed that nearly 70% of identified H&E-stained iNM granules fell within the threshold (Figure 4c, d). For extraneuronal pigments, we found that 90% of granules in all cases were smaller than 58.5 µm^2^ (Figure 4e, f). The distributions were again bimodal and showed that the majority of extraneuronal pigments were very small in size compared with iNM granules.

### The intracellular area occupied by neuromelanin granules

While the intracellular area occupied by iNM granules did not appear to be different by eye (Figure 5a), unstained iNM granules occupied a significantly larger neuronal area compared to H&E-stained granules (Figure 5b), potentially due to differences in tissue processing. Further analysis of the same sections measured before and after staining confirmed a significant reduction in the neuronal area of iNM following H&E staining, consistent with a change due to tissue processing (supporting information Figure S4). While H&E-stained iNM granules occupied a greater area of the neurons with increasing age, this relationship was lost when assessing the area occupied by the less processed unstained iNM with age (Figure 5c, d). These results reveal that histological processing impacts the intracellular area occupied by iNM.

**Figure 5.**
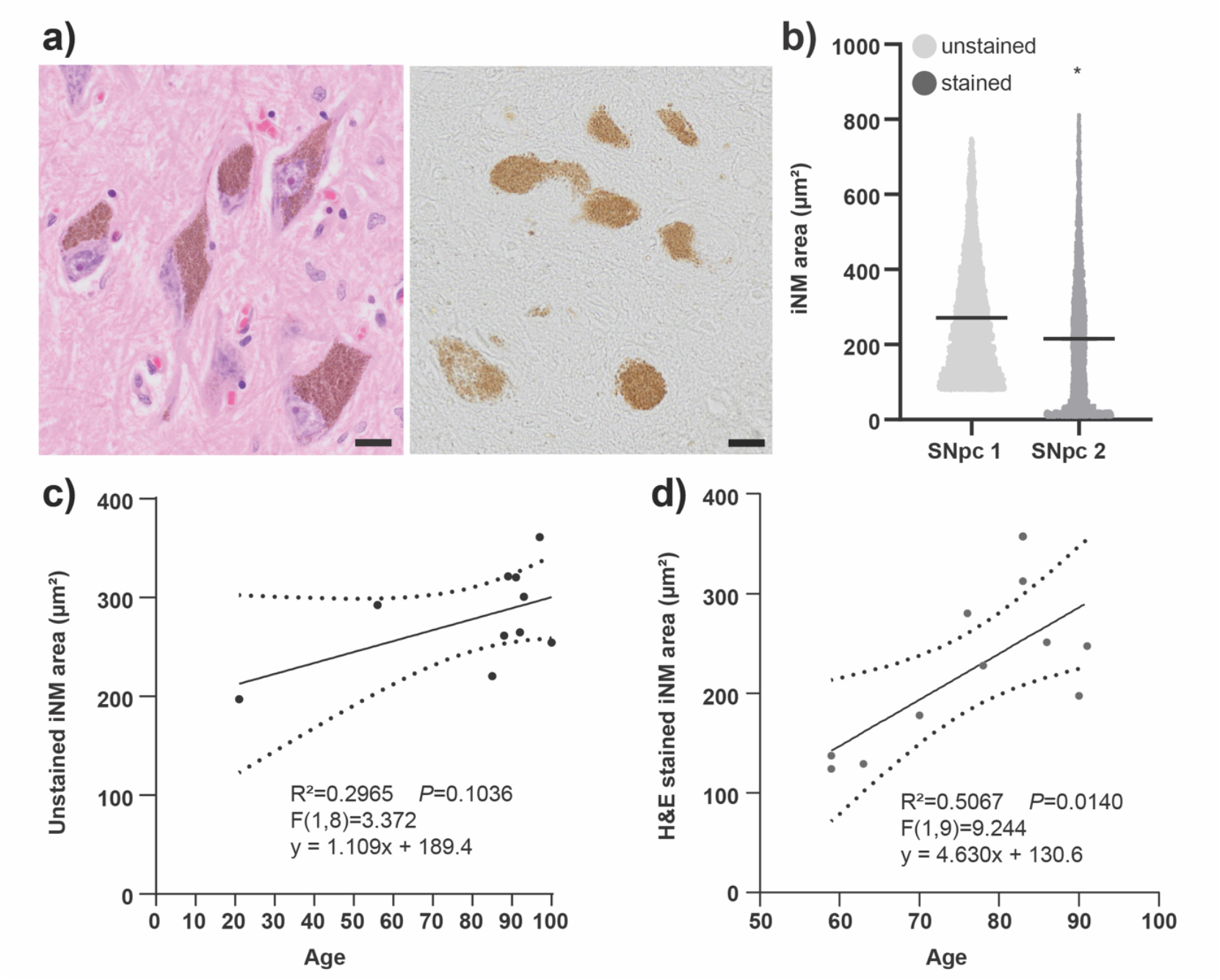
Analysis of the intracellular area occupied by neuromelanin. a) Representative images showing intracellular neuromelanin granules (iNM) granules in H&E-stained and unstained sections of the same regions of the substantia nigra pars compacta. Scale bars equal 20μm. b) Violin plots of the intracellular area occupied by iNM in the substantia nigra pars compacta (SNpc). The intracellular area occupied by unstained iNM was significantly larger than the intracellular area of H&E stained iNM, P < 0.0001, Mann-Whitney test. c, d) Scatterplots of the intracellular area occupied by unstained (c) versus H&E stained (d) iNM in healthy aging. There is an increase in the intracellular area occupied by iNM with increasing age which was significant in the H&E stained sections, linear regression P < 0.05.

### The optical density of intracellular neuromelanin

The optical density of iNM can be determined by its grey intensity. This measurement captures iNM accumulation and compaction. In previously used software, this feature was automatically quantified as a greyscale parameter. However, the TruAI platform is more comprehensive because it shows individual red, green and blue pixel measurements. To compute the optical density of iNM, the standard weighted equation from the red, green and blue colour intensity measures was used to estimate greyscale intensity:

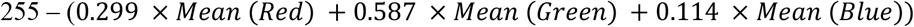

The estimated greyscale intensity did not significantly differ between unstained and H&E-stained iNM (supporting information Figure S3a). As expected, there is a significant increase in iNM estimated greyscale intensity with ageing (supporting information Figure S3b).

### Colour analysis of intracellular neuromelanin

Different types of peripheral melanins have different colours, so the analysis of colour could indicate similarities to other types of melanins [15]. We conducted a novel iNM colour analysis using TruAI red, green and blue pixel intensities. Initially, we evaluated the mean red, green and blue colour in unstained and stained sections. Notably, red, green and blue mean colour intensities showed significant differences in unstained and stained iNM granules, with the H&E stained sections having redder and bluer colour and less green colour (P < 0.0001 Mann-Whitney test, Figure 6a1-3).

**Figure 6.**
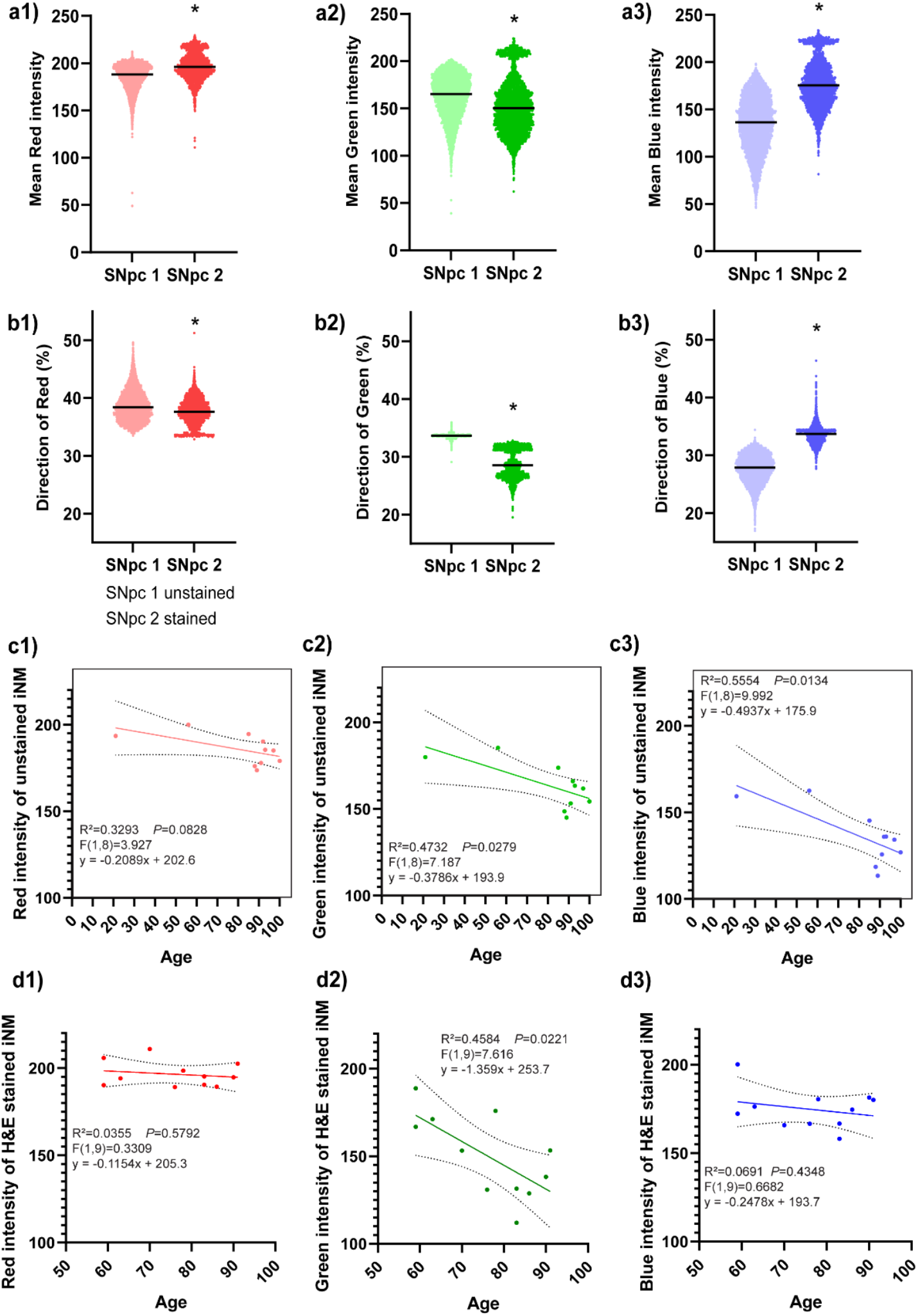
Colour intensity of intracellular neuromelanin granules (iNM). a1, a2, a3) Violin plots of the mean red, green, and blue intensities in unstained and H&E stained iNM granules in the substanita nigra pars compacta (SNpc). The H&E stained sections had more red and blue iNM compared to unstained sections, P < 0.0001, Mann-Whitney test. b1, b2, b3) Violin plots of weighted average individual colour contribution to iNM granules in unstained and H&E stained sctions showing that blue contributes most to the change in iNM colour in H&E sections, P < 0.0001, Mann-Whitney test. c1, c2, c3) Simple linear regression of red, green and blue intensities in unstained iNM showing significant reductions in gren and blue intensity with age, P < 0.05. d1, d2, d3) Simple linear regression of red, green and blue intensities in H&E stained iNM showing that only the green intensity reduces with age, P < 0.05.

To determine the individual colour contribution in iNM granules, we calculated the weight of each red, green and blue colour using the following equation:

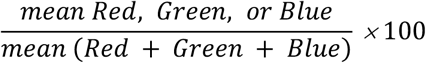

The individual colour contributions were also significantly different (P < 0.0001 Mann- Whitney test, Figure 6b1-3). Consistent with the absolute intensity, there was significantly more blue and less green in H&E-stained iNM granules (Figure 6b2-3), but now slightly less red colour in the H&E-stained iNM granules compared to unstained granules (Figure 6b1). This data suggests that the hematoxylin contributes most to the colour change in iNM.

Given that combined red, green and blue greyscale intensity increased with age, we were interested in seeing if any individual colour contributed to this change over time. Green intensity was significantly reduced in both unstained and stained iNM granules, whereas blue was only significantly reduced in unstained sections (Figure 6c1-3, d1-3). In contrast, red intensity remained stable in iNM granules under both conditions.

## Discussion

The most vulnerable neurons to PD are those that make the largely human-unique iNM. Understanding the role of neuromelanin in PD requires the standardisation of analysis platforms. We developed and validated an automated quantification workflow for reproducibly analysing neuromelanin in human post-mortem tissue sections. The protocol was tested on both unstained and H&E-stained tissue samples, allowing for the differentiation of iNM and extraneuronal pigments. In unstained sections, the TruAI software was trained to detect pigments using deep learning, followed by k-means clustering to establish iNM and extraneuronal pigment size thresholds. In contrast, nuclear staining allowed for manual characterisation of iNM and extraneuronal pigments in H&E-stained sections [16]. The system was then used to analyse parameters such as granule size, area occupied by iNM, optical density, and pigment colour. Our findings show that there is a substantial size difference between iNM and extraneuronal pigments, and that H&E staining affects the area occupied by iNM (through longer tissue processing).

Comparison of unstained versus stained neuromelanin granule size found two clusters in unstained sections when applying a k-means clustering size threshold. In H&E-stained tissue sections, this threshold was less sensitive for iNM granules, with nearly half of the iNM being smaller than the threshold in some cases. This exclusion of smaller stained particles in cells in H&E sections can be attributed to the similar dense blue staining of other smaller organelles, like ribosomes and mitochondria, as well as more shrinkage of the sections during the staining process [17-19]. The increased susceptibility of iNM granules to tissue shrinkage is attributed to lipid loss during dehydration and clearing [20]. Lipid droplets are key components of iNM, and the removal of these structures contributes to granule shrinkage [12, 21, 22]. This is supported by the observation that the iNM area was significantly smaller in neurons of H&E-stained tissue sections than in unstained neurons. Despite these artefacts, unstained and H&E-stained extraneuronal pigments were significantly smaller than the size of the iNM granules.

Histological processing also affected the intracellular area occupied by iNM. Consistent with previous findings, the intracellular area occupied by unstained iNM showed stability with increasing age [7]. However, the intracellular area occupied by H&E-stained iNM significantly increased with age. This has also been shown previously with the suggestion that the elderly have fewer neurons, but that remaining neurons are larger in size with no substantial change in the total perikaryon volume [23]. If such an age-associated change is related to tissue processing rather than actual neuronal size, the increased shrinkage of younger versus older brain tissue could suggest significant tissue composition differences with age. Recent analyses show a decreased volume and increased iron content in the SNpc with age [22, 24-26], with the decreased volume potentially suggesting shrinkage over time and increased iron potentially buttressing cell structures. This suggests that cell structures, and particularly iNM granules, may be less prone to tissue processing shrinkage in older individuals. In healthy ageing, NM granules sequester and store metals and cytotoxic quinones in lysosomes [22, 25, 26]. To maintain the stability of these pigments, lipid droplets support lysosomes through N-glycosylation [27]. Consequently, the older lipid droplets in iNM may exhibit chemical differences and be less susceptible to shrinkage during histological processing. In contrast, the estimated optical density showed the expected age-related linear increase [7], supporting the concept that increased iron may buttress the structure of older iNM. These measurements confirm the reliability of TruAI quantitation across differently stained tissue sections.

TruAI enabled new measurements of iNM, particularly colour analysis that is thought to be of functional relevance. Peripheral melanin consists of brown-black eumelanin and red-yellow pheomelanin [15], which have different functions - eumelanin acts as an antioxidant, whereas pheomelanin is a pro-oxidant [28]. The standardised TruAI quantification method explored these colour features in nigral neuromelanin pigments with and without H&E staining. The levels of red, green, and blue varied between unstained and H&E-stained iNM granules. H&E-stained granules showed higher levels of blue, likely due to the blue-purple haematoxylin pigment used in the staining process. Despite these differences, red was the predominant colour component in both types of iNM granules, indicating the presence of pheomelanin in healthy ageing. However, since red is just one colour, further evaluation of the combination of red, green and blue is needed.

Finally, we examined age-related colour changes. The red iNM component does not change with age, whereas green decreases significantly. Blue decreases significantly only in unstained iNM granules. In contrast, blue remained stable in H&E stained iNM granules, likely due to the presence of blue-purple haematoxylin pigment used in H&E staining that masks the true colour change. These combined colour alterations may indicate an increase in the brown-black eumelanin component. According to the additive red, green and blue colour model, vibrant red and green combine to form yellow, whereas a reduction in green intensity contributes to brown. The balance between yellow and brown pigmentation in peripheral melanin is controlled by differential gene expression in Drosophila. Specifically, a lighter-yellow peripheral pigmentation results from a mutation of the yellow gene [29]. In wild-type Drosophila, this gene contributes to dark brown eumelanin formation [30, 31]. This suggests that the loss of yellow may be related to the upregulation of antioxidant eumelanin in iNM in healthy ageing.

Overall, we have developed a standardised automated platform for the objective measurement of neuromelanin granules in tissue sections. iNM measurements were performed in two laboratories on different continents, giving similar results despite different methods, showing that the automated platform is accurate, easy to implement and reproducible. The TruAI platform allowed the accurate and standardised measurement of small intracellular granules that have been previously hard to measure and analyse. We found considerable differences in the size of iNM versus extraneuronal pigments and analysed the colour/type of these iNM granules with age, showing a change in colour, suggesting an increase in antioxidant eumelanin over time. Analysis of H&E-stained iNM granules supports a change in tissue and iNM components with advancing age. This standardised platform can now be used to determine disease and other associated changes in iNM that decrease the ability of catecholaminergic neurons to maintain their neuromelanin content prior to their neurodegeneration.

## List of abbreviations

AI: artificial intelligence
ROI: region of interest
H&E: Haematoxylin and Eosin
SND: substantia nigra pars compacta, dorsal tier
iNM: intracellular neuromelanin
SNL: substantia nigra pars compacta, lateral part
NM: neuromelanin
SNpc: substantia nigra pars compacta
PD: Parkinson’s disease
SNV: substantia nigra pars compacta, ventral tier

## Acknowledgements

The authors acknowledge: Microscopy Australia at the Sydney Microscopy and Microanalysis facility, University of Sydney, enabled by NCRIS; the Sydney Brain Bank in Australia, the Neurological Tissue BioBank at *IDIBAPS-Hospital Clínic* Barcelona and Biobank at IMIB-Instituto Murciano de Investigación Biosanitaria Pascual Parrilla Murcia in Spain, for human sample procurement; the ICTS-NANBIOSIS, U20/FVPR at VHIR, for assistance in histological processing; Heidi Cartwright for assistance with the figures.

## Author contributions

Conceptualisation and study design: Anastasia Filimontseva, Thais Cuadros, Zac Chatterton, YuHong Fu, Miquel Vila and Glenda Halliday. Data acquisition: Anastasia Filimontseva, Thais Cuadros, Amy Burke, Anahid Ansari Mahabadian and Zac Chatterton. Data analysis and interpretation: Anastasia Filimontseva, Thais Cuadros, Zac Chatterton, Amy Burke, YuHong Fu, Miquel Vila and Glenda Halliday. Reagent/materials/analysis tools: Anastasia Filimontseva, Thais Cuadros, Zac Chatterton, Amy Burke, YuHong Fu, Miquel Vila and Glenda Halliday. Drafting of the manuscript: Anastasia Filimontseva, Thais Cuadros and Glenda Halliday. Editing of the manuscript: Anastasia Filimontseva, Thais Cuadros, Zac Chatterton, YuHong Fu, Miquel Vila and Glenda Halliday. All authors have read and approved the final manuscript.

