## Supporting information for "Standardised TruAI automated quantification of intracellular neuromelanin granules in human brain tissue sections"

### Supporting information for Filimontseva *et al.* TruAI automated quantification of intracellular neuromelanin granules in human brain tissue sections

#Corresponding author: Professor Glenda Halliday, University of Sydney Brain and Mind Centre, 97 Church Street, Camperdown, NSW 2050 AUSTRALIA,, Tel: +61 2 9351 0888

**Table.** Information for the human brain material used in this study.

| **Brain Bank** | **MJFF code** | **Age** | **Sex#** | **ABC scores*** | **Analysis** |
|---|---|---|---|---|---|
| Sydney Brain Bank, Australia | Case 1 (control) | 21 | M | 0,0,0 | Unstained NM granules in figures 6,7,8,9 |
| Sydney Brain Bank, Australia | Case 2 (control) | 56 | F | 0,0,0 | Unstained NM granules in figures 6,7,8,9 |
| Biobank at IMIB, Spain | Control 9 | 59 | M | 0,0,0 | Stained NM granules in figures 6,7,8,9 |
| Biobank at IMIB, Spain | Control 11 | 59 | F | 0,0,0 | Stained NM granules in figures 6,7,8,9 |
| Sydney Brain Bank, Australia | Case 43 | 59 | M | 0,0,0 | Unstained NM granule size thresholding in figure 6 |
| Biobank at IMIB, Spain | Control 8 | 63 | M | 0,0,0 | Stained NM granules in figures 6,7,8,9 |
| Sydney Brain Bank, Australia | Case 41 | 69 | F | 0,0,0 | Unstained NM granule size thresholding in figure 6 |
| Sydney Brain Bank, Australia | Case 22 | 69 | M | 0,0,0 | Unstained NM granule size thresholding in figure 6 |
| Neurological Tissue BioBank at IDIBAPS-Hospital Clínic, Spain | Control 7 | 70 | F | 0,0,0 | Stained NM granules in figures 6,7,8,9 |
| Sydney Brain Bank, Australia | Case 19 | 72 | M | 1,0,0 | Unstained NM granule size thresholding in figure 6 |
| Sydney Brain Bank, Australia | Case 21 | 72 | M | 1,0,0 | Unstained NM granule size thresholding in figure 6 |
| Sydney Brain Bank, Australia | Case 23 | 72 | M | 0,0,0 | Unstained NM granule size thresholding in figure 6 |
| Sydney Brain Bank, Australia | Case 33 | 73 | M | 3,1,2 | Unstained NM granule size thresholding in figure 6 |
| Sydney Brain Bank, Australia | Case 18 | 75 | M | 0,2,0 | Unstained NM granule size thresholding in figure 6 |
| Neurological Tissue BioBank at IDIBAPS-Hospital Clínic, Spain | Control 1 | 76 | M | 0,0,0 | Stained NM granules in figures 6,7,8,9 |
| Sydney Brain Bank, Australia | Case 37 | 76 | M | 0,1,0 | Unstained NM granule size thresholding in figure 6 |

| | | | | | |
|---|---|---|---|---|---|
| Sydney Brain Bank, Australia | Case 27 | 77 | M | 0,1,0 | Unstained NM granule size thresholding in figure 6 |
| Sydney Brain Bank, Australia | Case 36 | 77 | M | 1,1,1 | Unstained NM granule size thresholding in figure 6 |
| Biobank at IMIB, Spain | Control 10 | 78 | M | 0,0,0 | Stained NM granules in figures 6,7,8,9 |
| Sydney Brain Bank, Australia | Case 20 | 78 | M | 0,0,0 | Unstained NM granule size thresholding in figure 6 |
| Sydney Brain Bank, Australia | Case 30 | 78 | M | 3,1,1 | Unstained NM granule size thresholding in figure 6 |
| Sydney Brain Bank, Australia | Case 32 | 81 | M | 0,1,0 | Unstained NM granule size thresholding in figure 6 |
| Sydney Brain Bank, Australia | Case 42 | 81 | M | 0,1,0 | Unstained NM granule size thresholding in figure 6 |
| Sydney Brain Bank, Australia | Case 24 | 82 | M | 2,0,1 | Unstained NM granule size thresholding in figure 6 |
| Sydney Brain Bank, Australia | Case 26 | 82 | M | 0,2,0 | Unstained NM granule size thresholding in figure 6 |
| Neurological Tissue BioBank at IDIBAPS-Hospital Clínic, Spain | Control 4 | 83 | M | 0,0,0 | Stained NM granules in figures 6,7,8,9 |
| Neurological Tissue BioBank at IDIBAPS-Hospital Clínic, Spain | Control 5 | 83 | F | 0,1,0 | Stained NM granules in figures 6,7,8,9 |
| Sydney Brain Bank, Australia | Case 25 | 84 | F | 2,1,2 | Unstained NM granule size thresholding in figure 6 |
| Sydney Brain Bank, Australia | Case 39 | 84 | M | 1,1,1 | Unstained NM granule size thresholding in figure 6 |
| Sydney Brain Bank, Australia | Case 3 (control) | 85 | F | 2,0,1 | Unstained NM granules in figures 6,7,8,9 |
| Neurological Tissue BioBank at IDIBAPS-Hospital Clínic, Spain | Control 3 | 86 | F | 0,0,0 | Stained NM granules in figures 6,7,8,9 |
| Sydney Brain Bank, Australia | Case 13 | 86 | F | 1,1,1 | Unstained NM granule size thresholding in figure 6 |
| Sydney Brain Bank, Australia | Case 5 (control) | 88 | F | 2,1,3 | Unstained NM granules in figures 6,7,8,9 |
| Sydney Brain Bank, Australia | Case 16 | 88 | M | 2,1,2 | Unstained NM granule size thresholding in figure 6 |
| Sydney Brain Bank, Australia | Case 35 (control) | 89 | M | 3,3,1 | Unstained NM granules in figures 6,7,8,9 |
| Neurological Tissue BioBank at IDIBAPS-Hospital Clínic, Spain | Control 2 | 90 | F | 0,0,0 | Stained NM granules in figures 6,7,8,9 |
| Sydney Brain Bank, Australia | Case 29 | 90 | M | 0,1,0 | Unstained NM granule size thresholding in figure 6 |
| Neurological Tissue BioBank at IDIBAPS-Hospital Clínic, Spain | Control 6 | 91 | F | 0,0,0 | Stained NM granules in figures 6,7,8,9 |
| Sydney Brain Bank, Australia | Case 11 (control) | 91 | F | 1,2,0 | Unstained NM granules in figures 6,7,8,9 |
| Sydney Brain Bank, Australia | Case 34 (control) | 92 | M | 2,1,1 | Unstained NM granules in figures 6,7,8,9 |
| Sydney Brain Bank, Australia | Case 17 | 92 | F | 0,0,0 | Unstained NM granule size thresholding in figure 6 |
| Sydney Brain Bank, Australia | Case 12 (control) | 93 | F | 1,1,0 | Unstained NM granules in figures 6,7,8,9 |

| | | | | | |
|---|---|---|---|---|---|
| Sydney Brain Bank, Australia | Case 14 | 94 | F | 2,1,1 | Unstained NM granule size thresholding in figure 6 |
| Sydney Brain Bank, Australia | Case 15 | 96 | M | 2,2,3 | Unstained NM granule size thresholding in figure 6 |
| Sydney Brain Bank, Australia | Case 40 | 96 | M | 1,2,1 | Unstained NM granule size thresholding in figure 6 |
| Sydney Brain Bank, Australia | Case 7 (control) | 97 | M | 1,1,0 | Unstained NM granules in figures 6,7,8,9 |
| Sydney Brain Bank, Australia | Case 8 (control) | 100 | F | 0,1,0 | Unstained NM granules in figures 6,7,8,9 |

[#]M=male, F=female

*National Institute on Aging-Alzheimer's Association guidelines for the neuropathologic assessment of Alzheimer's disease. Hyman BT, Phelps CH, Beach TG, Bigio EH, Cairns NJ, Carrillo MC, Dickson DW, Duyckaerts C, Frosch MP, Masliah E, Mirra SS, Nelson PT, Schneider JA, Thal DR, Thies B, Trojanowski JQ, Vinters HV, Montine TJ. Alzheimers Dement. 2012 Jan;8(1):1-13.

*Step by step guide for image acquisition scan project*

To ensure acquisition consistency, a “neuromelanin brightfield” scan project was created with the following steps:

1. The brightfield observation type and method were selected. The overview scan was performed at a 2x objective (NA=0.06), and the exposure was set to 295.57 μs. The first slide was manually observed and focused. All subsequent slides used this z-position as the focus depth (approximately -70um to -85um) (supporting information Figure S1a).
2. In the detail tab, a 20x objective (NA=0.8) was selected. The z-plane was set as ‘Normal’, and the exposure settings remained unchanged (supporting information Figure S1b).
3. ‘Prefocus’ was selected as the ‘focus method’, and ‘normal’ was selected as the ‘focus position density’ (supporting information Figure S1c).
4. In the ‘Naming and Saving’ window, ‘multi-layer image’ was selected to ensure the overview and detailed scans were saved as one file (supporting information Figure S1d1). This scan project can now be saved and applied as a single or batch scan (supporting information Figure 1d2). After the overview scan is complete, the ‘scan areas’ and ‘focus map’ can be adjusted to optimise the quality of the 20x scan (supporting information Figure S1e). Scan times vary based on sample size, objective, and modality. For example, a single plane of a 15 mm × 15 mm brightfield tissue section at 20x takes approximately 1 minute and 45 seconds for overview, focusing, and detailed scanning.

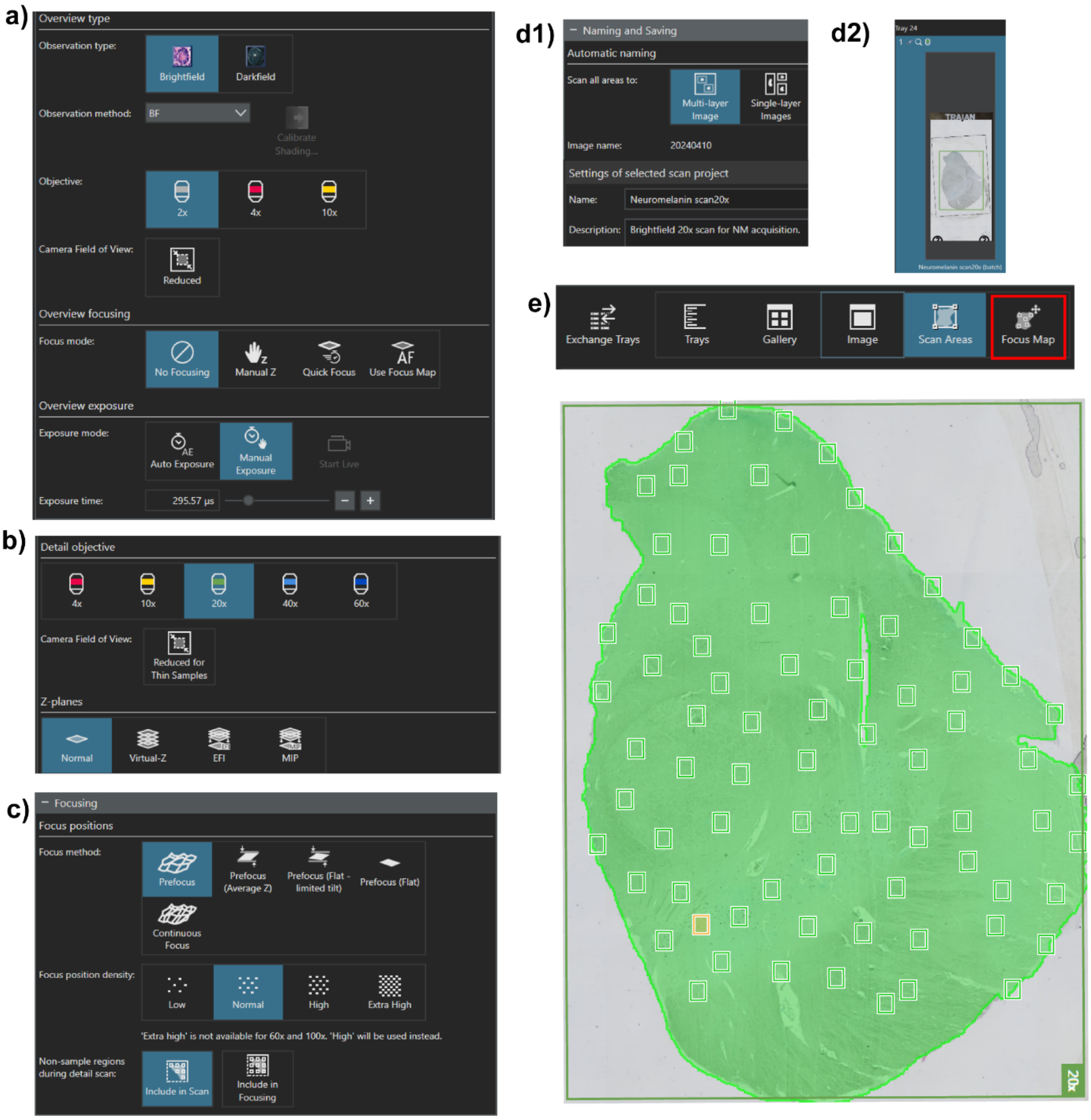


**Figure S1.** Standardised scan protocol for image acquisition. a) Imaging parameters for low-magnification overview. b) Selected objective and plane for detailed scan. c) Focusing parameters for higher magnification scan. d1, d2). Application of scan to sections. e) Creation of a focus map in the overview scanned section.

*Step by step guide for TruAI software training*

To train the software to automatically identify neuromelanin, two automated approaches were used. The first approach assessed unstained neuromelanin and the second approach assessed H&E- or cresyl violet-stained iNM and extraneuronal pigments. Neuromelanin training was performed as indicated below:

1. Open a scanned neuromelanin image in the Olympus VS200 desktop software (EVIDENT Technology GmbH, ver. 4.1.1 build 29408). In the 'Detect' tab in the top right-hand corner, select 'Training Labels' and create a new foreground class in level 1.
2. For unstained neuromelanin, name this class 'neuromelanin' and keep the 'Individual Objects' option unchecked (Figure 2a).
3. For H&E- or cresyl violet-stained neuromelanin three foreground training classes were created in level 1. These classes were 'iNM', 'extraneuronal pigments' and 'cells without pigment'.
4. It is essential to apply the same training label to each micrograph used for neural training. To ensure this, save the label to be used for neuromelanin training later (Figure 2b).
5. Select the automatically created 'background' class and draw a continuous area using the fill icon in the bottom left-hand corner under the 'Training Labels' section (Figure 2c).
6. Click on the 'neuromelanin' class and draw each neuromelanin granule in the highlighted background area. The thickness of the fill tool can be minimised to accurately outline the granules. Every neuromelanin granule must be drawn in the selected area (Figure 2d).

7. Apply steps 1-6 to 5-10 scanned sections and save them as training label images. These images were used for TruAI deep learning.

a)
b)
c)
d)

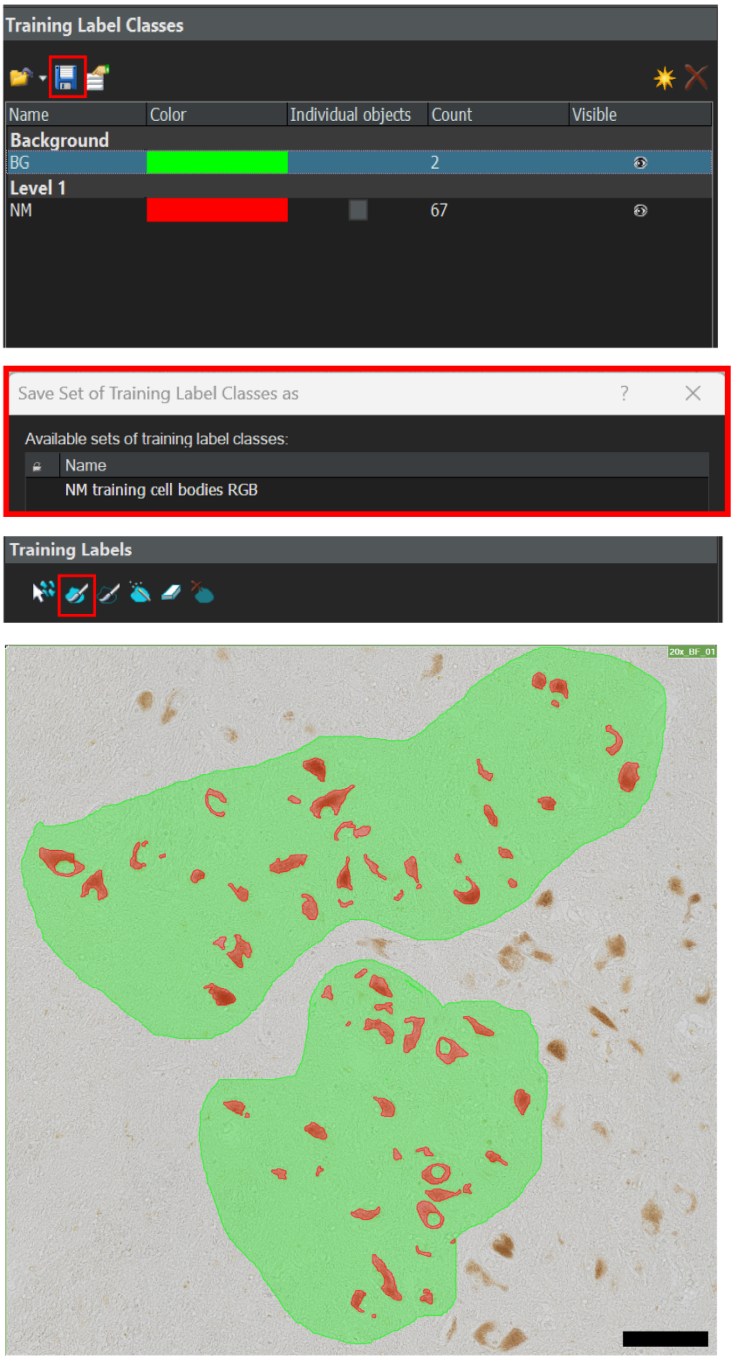


**Figure 2.** Training labels for neuromelanin (NM) quantification. a) Creation of a new training label class for NM granules. b) This training label must be saved and applied to sections. c) NM granules need to be accurately traced with the brush tool. d) Annotations need to be applied to 5-10 sections. Scale bar equals 100 µm.

*Step by step guide for TruAI deep learning automation*

Once at least five sections are annotated, the deep learning training can begin.

1. In the 'Deep Learning' tab, click 'New Training' and select the 'Image Segmentation' option. In the 'New Training: Input and Output' pop-up window, name the session 'Neuromelanin Neural Training' and load the sections with training labels by clicking the plus icon under the 'Images' section. Ensure that 'basic mode' is selected, and the input channel is red, green and blue. The individual objects option under training label classes should be left unchecked (supporting information Figure S2a).

2. In the next window select 'Specific Network (red, green and blue)' (supporting information Figure S2b). The required accuracy for automating neuromelanin quantitation is at least 0.85 similarity. For the first attempt, 'no limit' can be selected under training duration to allow for any modifications during the neural training.

3. Click 'Start' to commence training.

4. To increase the accuracy of neural training algorithm, more sections should be annotated (refer to TruAI software training above).

5. Select the most successful training algorithm by choosing the checkpoint with the highest similarity. Name this algorithm, provide a brief description, and save this neural network algorithm (supporting information Figure S2c). This algorithm will be used for all future automated quantification of neuromelanin.

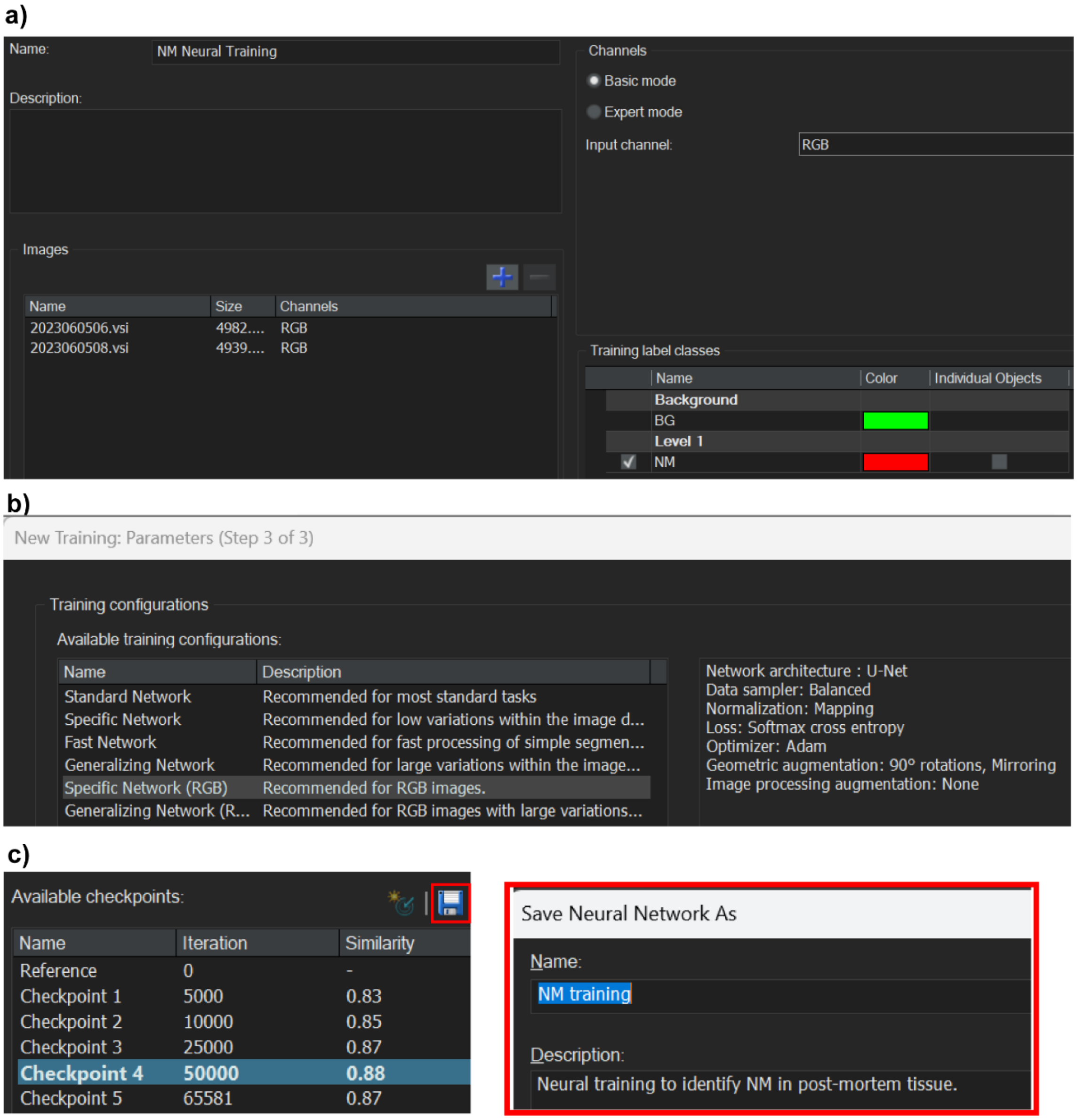


**Figure S2.** TruAI training on neuromelanin annotated sections. a) Neural training window interface. b) ‘Specific Network (red, green and blue)’ training configuration is suitable for automated neuromelanin identification. c) Neural training >0.85 similarity is required for reliable quantification. Once saved, successful training can be used for automated quantification.

*Step by step guide for fully automated TruAI quantification*

This neural network algorithm can now be applied to all scanned images for automated quantification of neuromelanin.

1. In the ‘Detect’ window of a scanned section, select the ‘Count and Measure’ tab in the bottom left corner. Click on the ‘Count and Measure’ drop-down menu (Figure 3a1) and select ‘New ROI’ to draw regions on the sections (Figure 3a2).

2. Once all ROIs have been completed, click ‘Neural Network Segmentation’ (Figure 3b1). Load the saved neuromelanin neural network and proceed to ‘Count and Measure on ROI’. Click on the ‘Dimension Selector’ tab to ensure this quantification is performed on the brightfield scan layer. The identified neuromelanin will be highlighted on the image (Figure 3b2).

3. The results for each individual neuromelanin granule will appear in the ‘Count and Measure Results’ (Figure 3c1). The parameters of interest can be chosen by clicking ‘Select Object Measurements’. For analysing iNM, size, intracellular area occupied by iNM and mean red, green and blue intensities can be selected from the drop-down menu (Figure 3c2).

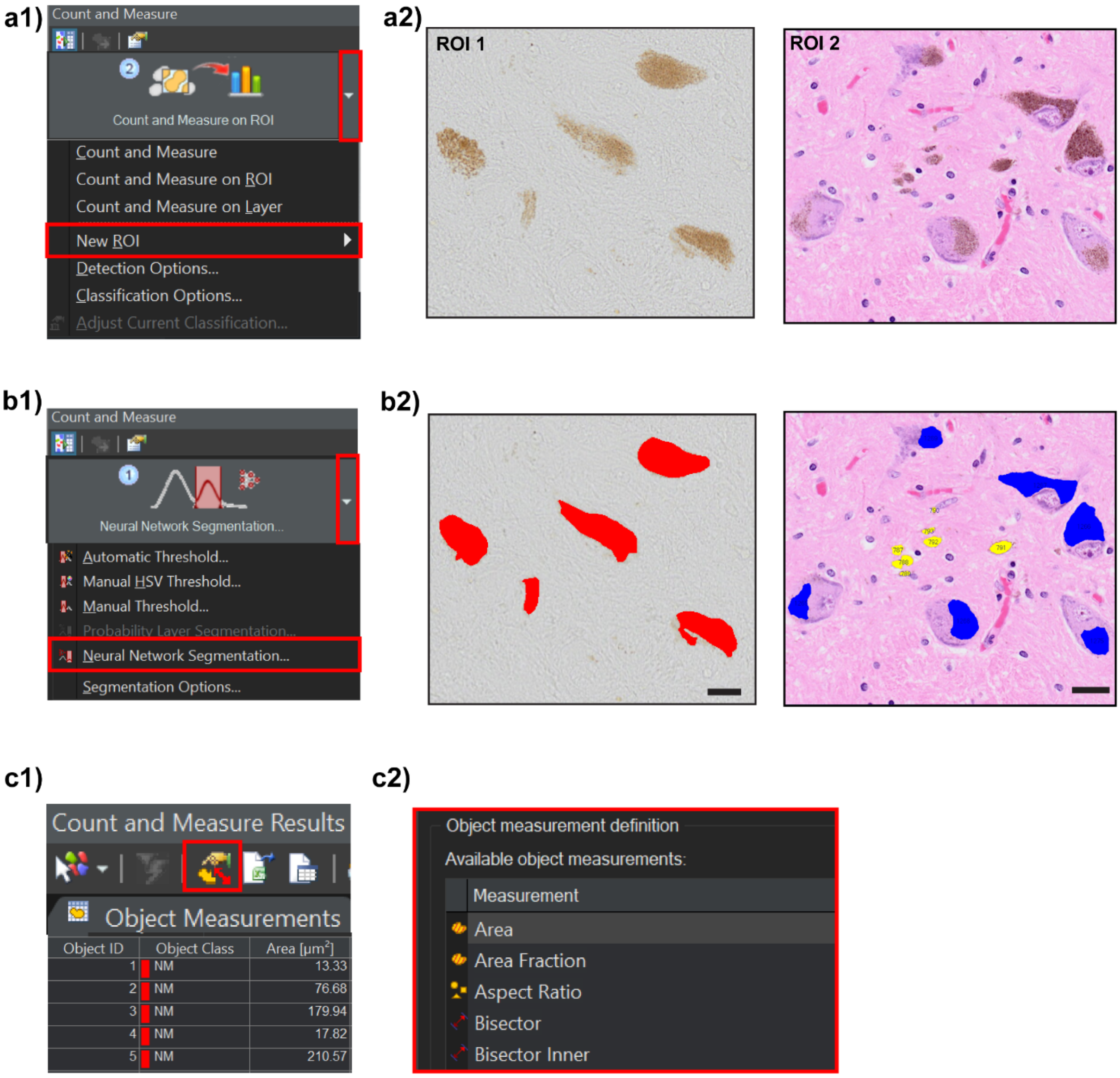


**Figure 3.** Automated TruAI quantification of neuromelanin. a1, a2) In selected sections, a new ROI needs to be drawn. b1, b2) The automated quantification of neuromelanin will run on the drawn ROI using the 'Neural Network Segmentation' option. Scale bars equal 20µm. c1, c2) A variety of features can be explored after completing neural network segmentation.

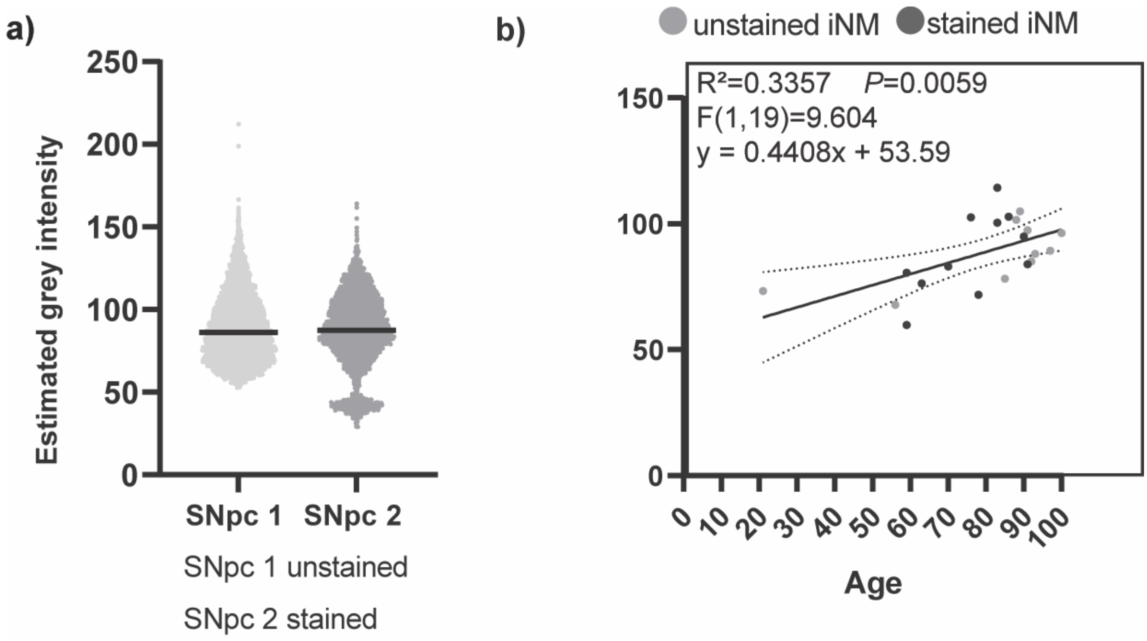


**Figure S3.** Estimated grey intensity for intracellular neuromelanin (iNM) in the substantia nigra pars compacta (SNpc). a) Violin plots of the estimated grey intensity for unstained and H&E stained iNM granules. b) Scatterplot of the average estimated grey intensity for unstained and H&E stained iNM during healthy aging. iNM optical density significantly increases with age, $P < 0.01$.

**Paired analysis of unstained versus stained neurons in a subset of cases**

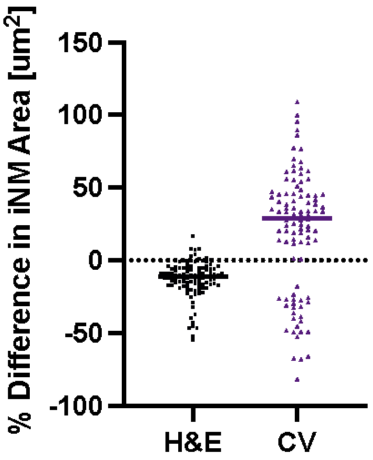


| | H&E | CV |
|---|---|---|
| De-paraffin | Same procedure was applied | |
| Rehydration | Same procedure was applied | |
| Staining | Haematoxylin & eosin | 0.5% cresyl violet |
| Dehydration | 70% ethanol 3 mins; 95% ethanol 3 mins; 100% ethanol 3 mins x 2; Xylene 3 mins x 2 | 100% ethanol 3 mins x 2; Xylene 3 mins x 2 |

**Figure S4.** Change in iNM area measured in single neurons first in unstained (dashed line) and then the same stained sections using two chemical staining methods and a comparison table of the methods. There was a significant reduction in the iNM area in neurons stained with H&E (paired t test t=10.89, df=104, $p<0.0001$, $R^2=0.5328$, r=0.9734, $p<0.0001$), while there was high variation with CV with some indication that CV stained a previously unseen iNM component in some neurons. The main difference in the methods is the increased dehydration of the tissue using H&E staining.
